# NOODAI: A webserver for network-oriented multi-omics data analysis and integration pipeline

**DOI:** 10.1101/2024.11.08.622488

**Authors:** Tiberiu Totu, Mattia Tomasoni, Hella Anna Bolck, Marija Buljan

## Abstract

Omics profiling has proven of great use for unbiased and comprehensive identification of key features that define biological phenotypes and underlie medical conditions. While each omics profile assists characterization of specific molecular components relevant for the studied phenotype, their joint evaluation can offer deeper insights into the overall mechanistic functioning of biological systems. Here, we introduce an approach where starting from representative traits (e.g., differentially expressed elements) obtained for each omics profile, we construct and analyze joint interaction networks. The resulting networks rely on the existing knowledge of confident interactions among biological entities. We use these maps to identify and describe central elements, which connect multiple entities characteristic for the studied phenotypes and we leverage MONET network decomposition tool in order to highlight functionally connected network modules. In order to enable broad usage of this approach, we developed the NOODAI software platform, which enables integrative omics analysis through a user-friendly interface. The analysis outcomes are presented both as raw output tables as well as high-quality summary plots and written reports. Since the MONET tool enables the use of algorithms with strong performance in identifying disease-relevant modules, the NOODAI software platform can be of a high value for the analysis of clinical multi-omics datasets.

**Availability and Implementation:** The platform is available as a web application freely accessible at https://omics-oracle.com. The source code is freely available from GitHub under the GPL3 license at: https://github.com/TotuTiberiu/NOODAI.

## Introduction

Omics characterization of biological samples allows for comprehensive identification and quantification of molecules present in the cells. Measurements of genome mutations, epigenome changes, as well as abundances of different forms of transcripts, proteins and metabolites, have proved to be of great use in understanding cellular processes that define biological phenotypes. Due to limitations in existing analytical methods, datasets obtained from different omics profiles are typically analyzed and interpreted individually. Nevertheless, due to strong interconnections among omics layers and the interplay of different types of molecules in exerting biological functions, a simultaneous evaluation of multiple omics profiles can allow for the identification of novel features representative of the system and critical to its function. Illustrations for this are phosphorylation and activation of transcription factors (TFs) that regulate downstream gene expression changes (Weidemüller, et al., 2021) or changes in metabolite levels that regulate protein degradation rates (Mitch, et al., 1994). Some omics profiles, such as proteomics, transcriptomics, and genomics, can be linked and analyzed together in a straightforward manner, while others, like epigenomics and metabolomics, require special considerations during the integration process (Babu and Snyder, 2023).

Several methods for integrating multiple omics profiles have been proposed in the literature, with some of them focusing on improving the interpretation of results obtained from the analysis of individual omics profiles and others on enabling the integration of different omics profiles into a unified framework (Picard, et al., 2021; Subramanian, et al., 2020). Among these, MOFA (Argelaguet, et al., 2018) and mixOmics (Rohart, et al., 2017) are prominent examples of statistical frameworks that integrate normalized measurements from different omics layers and jointly analyze them with the aim of identifying feature combinations (such as gene mutation, transcript and protein levels) that capture a high fraction of variation in molecular phenotypes. MOFA and mixOmics employ latent variable extraction through factor analysis or regression analysis, respectively. Other frequently used omics integration methods are WGCNA (Langfelder and Horvath, 2008), which is based on the usage of correlation analysis for data-driven network generation, BCC (Lock and Dunson, 2013) which leverages a Bayesian framework for consensus clustering across multi-omics datasets and MCFA (Brown, et al., 2023), which extracts latent factors based on a probabilistic model that combines dimensionality reduction with unsupervised learning to capture shared variations. Finally, iClusterPlus (Mo, et al., 2013) merges multiple omics datasets through a penalized latent variable regression model approach, while SNF (Wang, et al., 2014) represents a network-based integration approach that constructs sample-based similarity networks for each omics type and iteratively fuses them together until convergence towards a final clustering structure. While of great value, these tools often require specialized knowledge for interpreting the reported trends as well as significant programming skills for using the respective software tools.

To enhance the accessibility of methods for the integration of different omics layers and make them more user-friendly, a few web platforms have been developed. The 3Omics platform was one of the first web services to provide an easily accessible omics integration framework. Based on the user-provided features of interest from proteomics, transcriptomics, and metabolomics datasets, the web service offers pathway enrichment and feature co-expression analysis (Kuo, et al., 2013). Examples of two other frequently used and user-friendly platforms are OmicsAnalyst (Zhou, et al., 2021) and OmicsNet2.0-(Zhou, et al., 2022). These webtools utilize network analysis: OmicsNet relies on previously known interactions for building networks and OmicsAnalyst builds networks based on the inference analysis from the provided multi-omics features, such as for instance through correlation analysis, multi-view clustering and dimensionality reduction methods. While OmicsAnalyst has built-in functionalities for enabling pre-processing and data-driven integration of proteomics, transcriptomics, metabolomics and miRNAs raw measurements, OmicsNet requires input lists composed of user-selected elements that represent the respective types of omics layers in the studied conditions. OmicsNet also supports inclusion of information on significant single nucleotide polymorphisms and specific liquid chromatography-mass spectrometry peaks. Both platforms offer easy to use visualization tools and methods for functional enrichment analyses. Examples of other webtools for multi-omics analyses include Omics Integrator (Tuncbag, et al., 2016), GeneTrail2 (Stöckel, et al., 2016), MAINE (Gruca, et al., 2022), Mergeomics 2.0 (Ding, et al., 2021) and PaintOmics4 (Liu, et al., 2022). These tools frequently leverage interaction-based knowledge for both pathway enrichment analysis and data visualization methods. Results obtained from methods based on mathematical frameworks often pose interpretation challenges, and network-based approaches frequently use common graph theory methods that may not be optimized for all types of biological networks.

While extensive work has been undertaken to address the robustness and precision of different methods for the integration of omics layers, due to distinct properties and different distributions of measured data as well as unevenly distributed missing values, extracting clinically relevant outcomes from multi-omics profiles still presents a considerable challenge. Several of the existing methods exploit known and inferred connections among the biological entities and use graph theory tools for extracting biologically relevant information. The most often applied approaches are node centrality metrics and network decomposition tools, such as network modularization. A recent community-based Disease Module Identification DREAM Challenge (Choobdar, et al., 2019) has applied 75 network modularization methods on biological networks and comprehensively assessed their performance in the identification of closely connected disease-relevant network modules. The challenge assessed the performance of modularization tools on both single and multi-omics input datasets. The top three performing network modularization methods can be accessed through the MONET toolbox (Tomasoni, et al., 2020), which till now has been available as a container, but not as a webtool.

Here, we describe the Network-Oriented multi-Omics Data Analysis and Integration (NOODAI) webtool, an online platform for the combined analysis of multiple omics profiles. The tool takes as input user-provided lists of hits for different analyzed omics layers and maps them onto a high-confidence molecular interaction network, primarily based on known protein-protein interactions (PPIs). Through a user-friendly interface, NOODAI provides easy access to the top-performing modularization methods included in the MONET tool, as well as selected network centrality metrics. Due to its design, NOODAI can be used already with small sample sizes and few omics profiles. By performing pathway enrichment analysis on the MONET-identified network modules, the webtool highlights parallel signaling routes characteristic for each of the studied conditions. NOODAI additionally produces several publication-ready illustrations and a detailed report in which the top central features supported by independent input datasets are functionally described. The tool can be fully customized to study any combination of omics datasets across species as it provides a possibility to adapt all the underlying databases files. The required databases are pre-loaded on the server for human, mouse, rat and 10 other species.

## NOODAI analysis pipeline overview

NOODAI is a cloud-based web service that supports joint interpretation of features identified as significant in the analysis of omics datasets. The required inputs are lists of protein or small molecule identifiers (IDs), which should be provided individually for every omics layer (i.e. for genes and transcripts representative UniProt IDs should be used and for metabolites ChEBI IDs). The input lists should contain elements that characterize the samples in a studied condition and can be merged into an interaction network, such as significantly up- and down-regulated expression values. The lists of significant features for different studied conditions should be organized as separate sheets within individual Excel files, each omics layer should have a separate Excel file and Excel files for all studied omics layers must be compressed together into a single archive. NOODAI is not constrained by the number of omics profiles and studied conditions it can accept, being effective as a single-omics tool as well. The formatting requirements are described in the “Instructions” section of the platform and in the Supplementary materials. In addition, an example input dataset is provided on the website. NOODAI maps the provided elements onto biological networks, identifies elements that are central for network connectivity and reports network modules with highly connected elements, which we identify with the MONET tool. For each module, information is provided on whether its elements are enriched in specific signaling pathways. This provides a broader context on parallel biological processes associated with a studied condition. Finally, a summary report is generated: It includes information on the biological functions of the most central elements, it highlights central elements with regulatory roles, such as kinases and transcription factors, and it provides publication-ready illustrations; for instance, circular plots of most connected elements.

As a first step, the input lists with elements that are relevant for different omics layers (Figure 1**A**) are mapped to a knowledge-based PPI or protein-small molecule network. The readily available network includes pairwise interactions from STRING (Szklarczyk, et al., 2019), BioGrid (Oughtred, et al., 2021) and IntAct (Del Toro, et al., 2022) databases, filtered according to the respective database-defined criteria in order to include only high-confidence interaction pairs. It is possible for the user to provide their own reference interactions, but the NOODAI platform includes reference datasets with high-confidence interactions for 13 species. Interaction networks are first constructed for all individual omics layers separately and then merged through concatenation. The network with the highest number of members is used for further analysis (Figure 1**B**) and smaller disconnected networks are discarded.

**Figure 1.**
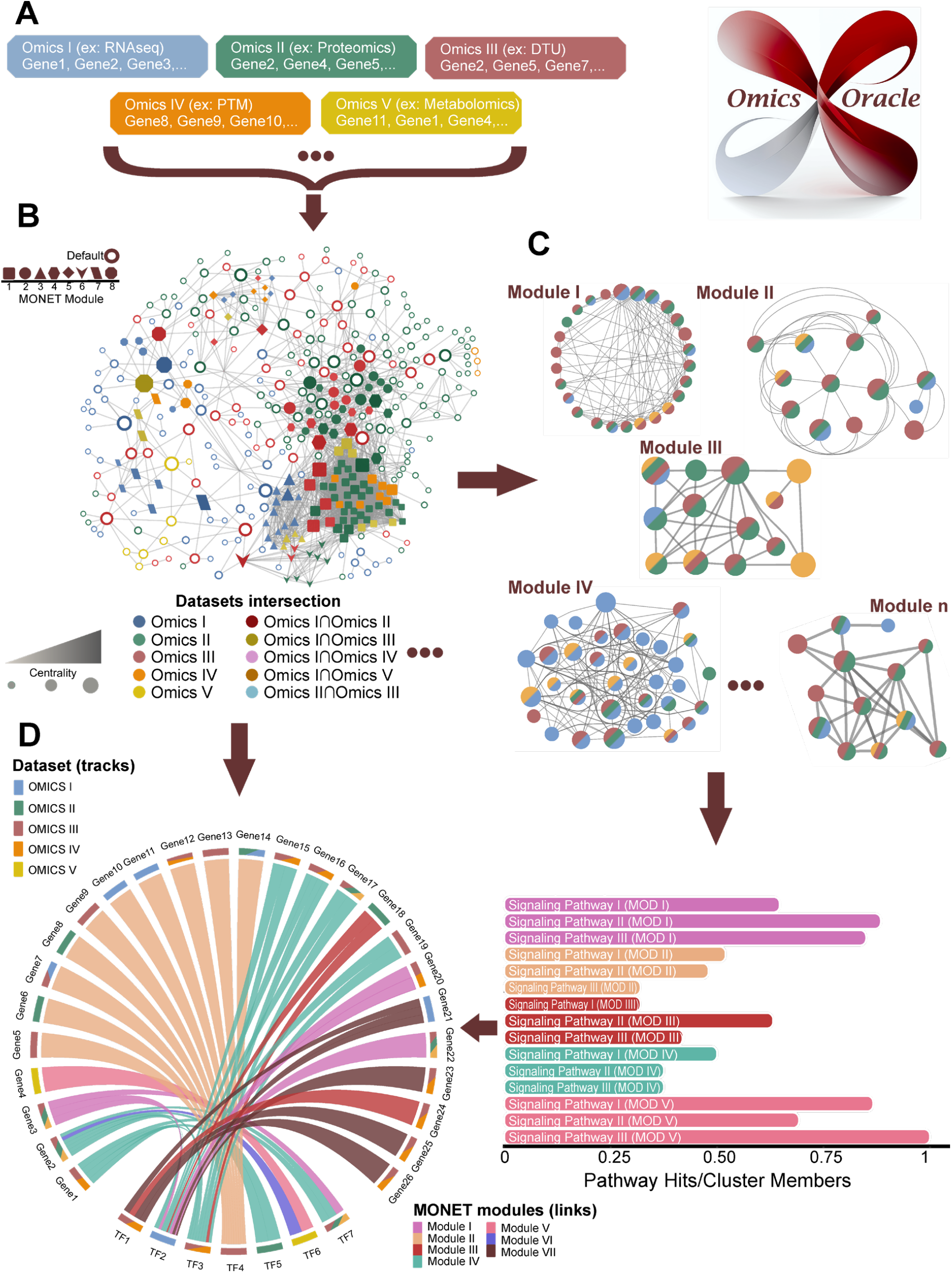
NOODAI analysis pipeline. (**A**) The platform takes as input lists of UniProt or ChEBI identifiers that are representative for each analyzed omics profile in a given condition. (**B**) For each condition, input lists from different omics layers are integrated into an interaction network. Following, the current-flow betweenness centrality metric is computed for each node, i.e. representative protein, in the network, (**C**) and the global network is decomposed into modules using the MONET tool. (**D**) The last step consists of generating summary-level plots. For each module, information is gathered on the pathways that its constituent elements are annotated with, statistical enrichment is assessed and fractions of elements annotated with the most strongly enriched pathways are shown as barplots. Afterwards, transcription factors with the highest centrality scores are shown together with their interaction partners which themselves had high centrality scores. Information on the shared modules and omics layers in which the proteins were highlighted is also included in the plot.

Following the network construction, NOODAI calculates network centralities to assess the importance of individual proteins for the overall network connectivity. Network centralities of the constituent nodes are calculated using the metrics provided by the CINNA R package (Ashtiani, et al., 2019), and are made available to the user as output. For the default metric used in the NOODAI pipeline, the Current-flow betweenness centrality was chosen. This centrality metric assesses the shortest paths that connect network elements through each respective protein, and additionally includes contributions from all possible paths by accounting for the information flow through random walks (Newman, 2005). In this way, nodes that represent network bottlenecks are effectively identified. Network metrics based on betweenness centrality were reported to be particularly suitable for highlighting crucial elements in regulatory biological networks (Alvarez-Ponce, et al., 2017; Yu, et al., 2007). Following, the network nodes are ranked according to their centrality scores. Elements that are within the top 15% nodes with the highest centrality and were additionally listed as significant in ¾ or more of the studied omics layers are referred to as robust nodes.

With the centrality scores determined, the joint network is then decomposed into modules in order to identify key communities of highly connected elements. In addition, the functional roles associated with individual modules are assessed. Non-overlapping network modules are determined using the MONET tool (Tomasoni, et al., 2020) (Figure 1**C**). The tool enables usage of three network modularization algorithms, which in the DREAM challenge were able to effectively single out disease-relevant functional modules. With the default settings, the NOODAI web platform uses the modularity optimization algorithm (M1) due to its lack of a stochastic component and its low computational requirements (Tomasoni, et al., 2020). The modularity optimization method has its foundation in a technique named Multiresolution and it evaluates the topological scales at which modules may be found (Arenas, et al., 2008). However, the user can also select any of the two other MONET methods, R1 or K1. Additionally, as a default, an undirected network is assumed and the desired average nodes degree in the identified MONET modules is set to 10. We expect that closely interacting proteins will often share functional roles and be involved in the same signaling pathways. Therefore, after the modules are defined, we assess whether proteins in the same module are enriched in any specific pathway. For this, we first note the ratio between the number of proteins in a MONET module associated with a specific pathway and the total number of elements in the module and then assess if the trend is statistically significant. A default over-representation analysis is performed using pathway annotations from the Reactome database (Milacic, et al., 2024), but the platform also supports the usage of several other databases with functional and pathway annotations. Pathway over-representation is assessed with the Fisher’s exact test and the correction for multiple testing is performed with the Benjamini-Hochberg’s test at the level of modules. Regardless of whether the statistical significance threshold was reached, the three most significant pathways from the five largest modules are visualized as a barchart (Figure 1**D**), which is included in the NOODAI summary output. Different MONET configuration parameters as well as pathway databases that are readily available for the enrichment analysis are described in more detail in Supplementary materials and in the “Instruction” section of the platform.

Given the central role of transcription factors (TFs) in regulating cellular processes, defining phenotypes, and guiding development, NOODAI focuses on identifying key TFs in the network. The most central TFs are highlighted on a circular diagram illustration together with their selected interaction partners, which themselves were ranked among the network elements with the highest centrality scores. This plot is included in the NOODAI summary output (Figure 1**D**). The color scheme in the circular illustration entails information on signaling pathways enriched in the modules to which the shown TFs belong.

The web platform includes a demo dataset that allows users to easily understand the data formatting and explore the default setting parameters. The dataset comprises lists for different omics layers with the most significant results obtained after analyzing proteomics, phosphoproteomics, transcriptomics, and splicing datasets from macrophages in different functional states. The lists constitute of most significantly upregulated and downregulated elements after pairwise comparisons of human primary macrophages in the *in vitro*-generated phenotypes known as M1, M2a, and M2c. Further information about the demo dataset can be found in the “Tutorial” and “Instructions” sections of the platform, as well as in the original publication of the datasets (Totu, et al., 2024).

Finally, to ensure a comprehensive overview of the main findings, a report including information on all central network entities (top 15%) and results of the network decomposition analysis is automatically generated. The selected nodes are listed together with information on the number of omics layers in which they were identified as significant, basic functional description and notion if the node is a known cellular regulator and functions as a TF or a kinase. If the node is a member of a module enriched in a specific signaling pathway, this information is also included in the report.

## Conclusions

NOODAI web service allows for the simultaneous evaluation of multiple omics profiles, and is able to capture system-level characteristics that underlie the studied biological phenotypes. NOODAI provides easy access to a network-based integrative method for the joint characterization of significant features from the analyses of multi-omics profiles. The method is based on the generation of knowledge-based high-confidence interaction networks, which are analyzed both at the level of individual elements as well as at the level of tightly linked network neighborhoods or modules. Individual elements are assessed according to their centrality in the network, reoccurrence as significant in different omics layers and their functional roles. Modules are further investigated for the enrichment in specific signaling pathways or functional roles. In addition to the reference resources and the default analysis settings available on the platform, the knowledge-based interaction database, reference functional annotations and analysis parameters can be tailored to the user‘s needs. This also makes the tool applicable across different species. Human, mouse, rat and ten other species have all the required databases pre-loaded. NOODAI differentiates itself from similar tools by enabling users to fully tailor their analysis parameters and databases, while offering access to biologically specific network decomposition methods through MONET. The platform, accessible at https://omics-oracle.com, delivers main results as raw data in a tabular format together with publication-ready summary-level plots and a comprehensive summary report. NOODAI supports the delivery of clinically relevant insights from the combined analysis of multiple omics profiles, without the need for user’s experience in programming and statistics.

## Supporting information

NOODAI_Supplementary

## Acknowledgments

The presented work greatly benefited from the feedback provided by the Particles-Biology Interaction team members.

## Conflict of interest

The authors declare no competing interests.

## Funding

This research was supported by the Uniscientia Foundation, Swiss Cancer Foundation and Empa-KSSG seed grants as well as Julius-Müller Stftung and Sassella Stiftung.

## Notes

https://omics-oracle.com/

### Competing Interest Statement

The authors have declared no competing interest.

