## Supplementary material for "NOODAI: A webserver for network-oriented multi-omics data analysis and integration pipeline": NOODAI_Supplementary

This supplementary material provides a detailed workflow of the NOODAI pipeline, the main webtool interface functionalities as well as technical validations for the default metric selections.

### Contents


I. NOODAI analysis pipeline

II. Case study with results validation

III. NOODAI workflow

#### 1. NOODAI analysis pipeline

The NOODAI software platform supports inference of the most important molecular features from the joint analysis of multiple omics layers. Its architecture relies on the usage of pre-existing interaction networks for the mapping of entities obtained through the pre-analysis of individual omics profiles. The input for the pipeline is represented by lists of features (i.e. proteins, genes, transcripts, small molecules) obtained from the analysis of one or more omics datasets, such as entities that are significantly up- or down-regulated in a specific phenotype or condition of interest. The set of features from one omics layer that characterizes a particular condition is referred to as a profile. The default configuration available on the web platform requires protein identifiers (IDs) for the input features (i.e. representative protein IDs for differentially expressed genes) or small molecule IDs. By using a different type of knowledge-based interaction network, the analysis pipeline can be extended to accommodate any other feature lists important for characterizing the condition or phenotype of interest. For a consistent analysis, it is crucial that the features derived from various omics datasets can be logically integrated into a cohesive interaction network. The NOODAI webtool output delivers a comprehensive characterization of the studied biological conditions, presented through summary plots, tables, and a detailed report. The key elements involve analyzing the constructed networks using centrality metrics and functional characterization of network modules through the application of the MONET network decomposition tool. The latter allows for the inference of parallel signaling routes that affect molecular entities in different omics layers.

##### 1.1 Protein-protein interaction network construction

After providing the pre-selected features' as UniProt or ChEBI IDs for each of the analyzed omics layers, the NOODAI maps individually the features from each omics profile to the pre-existing network. When using the default settings, a protein-protein interaction (PPI) network and a protein-small-molecule network are constructed. For this, high-confidence interaction pairs collected from public knowledge databases, which are readily available on the webserver, are queried and only those for which both elements are in the input feature list are kept. The interactions are collected from STRING (Szklarczyk, et al., 2019), BioGrid (Oughtred, et al., 2021) and IntAct (Del Toro, et al., 2022) databases. STRING, database version 11.5 was filtered to include only entries with a combined score above 0.7 (starting from the complete interactions data from all sources). From the BioGRID database version 4.4.218, a mitab data file was used and only physical interactions supported by independent validations and an associated confidence value were kept. For the high confidence interactions from IntACT, the default psimitab database from 13/07/2022 was filtered to keep only interactions with a confidence score above 0.7. The following thirteen NCBI organisms' taxonomies have pre-loaded reference interactions and database files on the server and can be directly queried with protein and small molecule entries: 9913, 6239, 9615, 7955, 44689, 7227, 9031, 10090, 10116, 4932, 4896, 8364, 9606. For other organisms, custom reference interaction files must be uploaded to the webtool by the user (see Section III).

After generating the networks for each omics profile, the networks are merged together through concatenation. All nodes and edges in the joint network are unique. The networks are undirected and unweighted. Reliance on known interactions often results in many smaller isolated networks built from a handful of elements. In NOODAI, only the network which includes the highest fraction of the provided entities is further analyzed.

##### 1.2 **Node's centralities**

After the construction of the PPI network, centrality scores are calculated for each node in the network. For this, we use the CINNA R package (Ashtiani, et al., 2019). The package computes 49 distinct centrality metrics and NOODAI reports 35 of them as some metrics demand excessive computational resources when scaling. When interpreting results for the PPI networks, which rely on the overlap of results from multi-omics analyses with known interactions, it is important to bear in mind possible confounding trends.

Firstly, public knowledge PPI databases on one hand, miss important interactions that are not yet validated and on the other hand, tend to have well-studied proteins over-represented due to the literature bias. Secondly, most network analysis methods from graph theory were not designed to convey information with biological meaning. NOODAI reports current-flow betweenness score as a default centrality measure. This metric is computed in addition to the ones found in the CINNA package. Previous work has shown that betweenness centrality performed better than degree of direct interactions (i.e. "hub-ness") in identifying central regulatory genes with significant biological relevance (Alvarez-Ponce, et al., 2017; Yu, et al., 2007). Current-flow betweenness accounts for additional routes of information flow in the networks. In section II a comparative study with the other centrality metrics is provided. By default, NOODAI determines the importance of a node based on both its centrality score and the number of omics evidence supporting its presence. Nodes that are found in at least half of the input lists and which are in the top 10% of most central nodes are categorized as 'salient nodes' and are deemed as likely important for the studied profiles.

##### 1.3 **Network decomposition**

After extracting the key features for each profile under study, additional functional characterization of the profiles is performed after the MONET decomposition of the PPI networks (Tomasoni, et al., 2020). MOENT supports usage of algorithms represented by the M1, R1 or K1 decomposition methods. The M1 method was chosen as the default option due to its scalability and reproducibility (Tomasoni, et al., 2020). By default, the edges are assumed to be undirected and the average desired node degree for the modules is set to 10. The MONET methods were evaluated specifically in the context of biological systems and represent the top-performing algorithms of the 'Disease Module Identification DREAM Challenge', a community effort to develop unsupervised network modularization algorithms for biological networks (Choobdar, et al., 2019). Network decomposition is applied to the joint PPI network and the identified modules of highly connected proteins will often contain members from multiple input omics layers. Proteins in the same module are expected to often be involved in the same biological processes and signaling pathways. In order to investigate which signaling pathways are associated with individual modules, annotations from the Reactome (Milacic, et al., 2024), Wikipathways (Agrawal, et al., 2024), BioCarta (Nishimura, 2001), PID (Schaefer, et al., 2009), NetPath (Kandasamy, et al., 2010), HumanCyc (Romero, et al., 2005), INOH (Yamamoto, et al., 2011) and SMPDB (Jewison, et al., 2014) signaling pathways databases are used and enrichment compared to the background of all proteins found in the full-scale PPI network is assessed. This analysis is conducted only on modules with 10 or more members. A module is assumed to be relevant for mapping independent functional routes that characterize the studied phenotype if at least 50% of its members are associated with a specific signaling pathway. This is particularly of interest when it helps to identify proteins that come from different omics profiles but are grouped together in the network and have shared functional relationships. Criteria for the required fraction of proteins in a module with a shared function can be adapted by the users. Additionally, the Benjamini-Hochberg (Benjamini and Hochberg, 1995) corrected p-values (FDRs) calculated for the statistical enrichment compared to the overall network are available and can be used as an additional filtering threshold. FDR thresholds are not applied by default. All pathways identified as shared among the majority of modules’ members are reported to allow for a comprehensive evaluation of the roles that the module can be associated with. However, for the final presentation of results, it is advisable to filter out pathways whose members and functions strongly overlap. Cumulative interpretation of representative functions performed by individual modules can be used to generate an overview into the main signaling flow axes that underlie the studied phenotype.

##### 1.4 **Summary plots and report generation**

Next, results generated in the analyses above are aggregated into summary-level plots and a comprehensive report. In addition, the edges in each of the MONET modules are extracted to support the visualization of modules using Cytoscape (Shannon, et al., 2003). An R script is provided for this (Totu, 2024). In the report, the top three signaling pathways (based on the FDR values) for the largest five modules are illustrated in the form of a barchart where the ratio between the number of pathway members and the total number of members in the respective module is plotted. It is important to note that these top 3 pathways can be highly overlapping and have almost identical members. The overlap between the pathways can be evaluated by assessing the composition of their members. This information is available in the Excel tables in the results folder. In addition, the report includes a circular representation diagram that highlights the most central transcription factors (TF) and their connecting proteins, which themselves have a high centrality score. The connection between each interacting protein and TF is colored based on the module in which the interacting protein is found. Only interacting proteins that are among the top 30% of central nodes in the full-size networks are considered during the generation of the plot, the maximum number of represented TFs is set to 7 and the maximum possible number of interacting proteins to 25. The barplot with pathways associated with each module and circular diagrams aim to offer the user a fast and comprehensive insight into the analyzed profiles. Besides this overview, the main results are presented in an automatically generated report in which the most central nodes are presented, and possible major signaling axes are highlighted. Information is also provided on whether the same feature was present in multiple input lists and nodes ranked among the top 15% of most central ones that are also found in at least 75% of the input omics lists are named as robust. Kinases found within the top 15% of central nodes are also reported for the analyzed profiles. Pathways for which 70% or more of the members share the same function are also prominently highlighted. The summary report contains information for all the characterized phenotypes or conditions with the circular and signaling pathways plots produced separately for each condition.

#### 2. Demo dataset and rationale behind default settings

Macrophages are immune cells that exhibit a high degree of phenotypic plasticity and are known to be involved in a number of pathological conditions (Totu, et al., 2024). In the context of cancer, macrophages are double-edge swords exhibiting both pro- and anti-tumorigenic phenotypes by either supporting or inhibiting anti-tumor inflammatory activity. Even though extensively studied, our current understanding of the main phenotypic drivers is largely confined to pro-inflammatory macrophages. Understanding the differences in the main signaling routes and phenotypic traits between different types of macrophages could impact the rational design of novel cancer therapeutics.

Here, we applied the NOODAI pipeline on previously generated omics datasets for human blood-derived pro-inflammatory *in vitro* M1 (LPS and IFN-γ stimulated) and *in vitro* immunosuppressive M2a (IL-4 and IL-13 stimulated) and M2c (IL-10)-macrophages (Totu, et al., 2024). The datasets included next-generation sequencing transcriptome measurements that indicated gene expression levels and allowed inference of splice isoforms as well as mass spectrometry-based proteomics and phosphoproteomics quantitative measurements. These datasets are available in a reduced form as Demo on the web platform. Due to resource management, the Demo dataset contains only 20% of the original omics data, but nonetheless preserves the original trends (Totu, et al., 2024). Throughout this section, we will follow each analysis step with direct application to the Demo data and present the rationale behind the default settings chosen for the NOODAI pipeline.

The analyzed omics profiles in M1, M2a and M2c macrophage states were represented by features (proteins and genes) that were found to be upregulated in one phenotype when directly compared to another. The criteria for this was a log2FC threshold of 1 and FDR threshold of 0.05 for (phospho-)proteomics and a log2FC 2 and FDR threshold 0.05 for transcriptomic data. Pairwise comparisons between M1 and each of the M2 states were performed on all available omics layers and significantly up- and down-regulated entities were noted separately. This resulted in the six input lists in the demo datasets for each analyzed layer, with the exception of the splicing layer which included only differentially used transcripts.

For each input list, i.e. significantly up or down-regulated entities, PPI networks are built for each omics layer and then a joint network is generated through concatenation. Next, current-flow betweenness centrality is calculated for each node within the networks. In case of the demo dataset, this highlighted proteins such as STAT1, NFKB2 and PML for pro-inflammatory M1 macrophages, and CSF1R and ITSN1 for M2 macrophages. Central regulatory roles of these proteins in the respective phenotypes are supported by numerous previous studies (Chen, et al., 2023; Ordentlich, 2021).

In order to assess in more detail the role of the nodes ranked in the top 10% of the current-flow betweenness centrality scores in underlying macrophages clinical phenotypes, we performed analysis with the VOSviewer bibliometric tool (Van Eck and Waltman, 2014). For this, all clinical studies on macrophages from Embase from 2012 onwards were screened using the search term: “((human OR primary) NEXT/3 macrophage*) AND [humans]/lim AND [clinical study]/lim)”. Only the terms with at least 3 mentions were selected. This showed that at least one tenth of the most central nodes in each contrast was mentioned in the clinical studies that involved macrophages. To evaluate NOODAI-computed centrality algorithms, the percentage of network proteins associated with clinical trials that fall within the top 10% of central nodes, relative to the total number of clinically associated network proteins, was used as a benchmark. On average, current-flow betweenness centrality aggregates 30% of the network proteins associated with clinical trials within the top 10% of central nodes (Figure S1). Eccentricity, Group and Topological Coefficient centralities show similar performance.

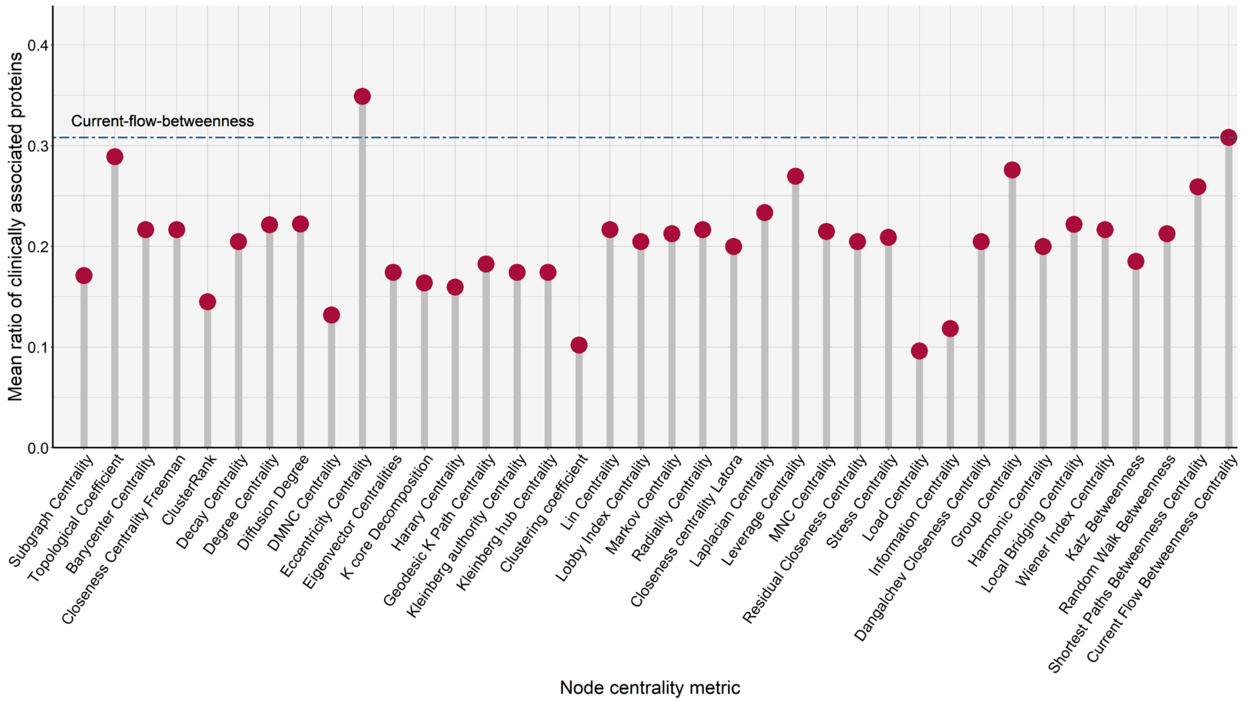


**Figure S1.** The average ratio of clinically associated proteins within the top 10% of central nodes, relative to the total number of clinical proteins found in the datasets, for each of the centrality metrics computed by NOODAI.

Following, the MONET network modularity optimization method (M1) is used for the network decomposition. For the resulting modules, we assess whether the majority of their members are associated with specific signaling routes using as a reference signaling pathways over-represented (FDR < 0.01) from 503 macrophage-specific genes annotated in the Human Protein Atlas that aggregates single-cell RNAseq studies from 14 different tissues (Karlsson, et al., 2021). We found that each module of interest (i.e., those with at least 50% of members associated with a pathway) contained at least one macrophage-specific signaling pathway. For M1 macrophages, the main parallel signaling routes included IFN-$\gamma$, Toll-like receptor signaling, NF-$\kappa$B signaling, NOD-like receptor signaling, TNF and JAK-STAT signaling. This convincingly recapitulated the known biology of pro-inflammatory macrophages (Chen, et al., 2023). The MONET modules associated with immunosuppressive M2 *in vitro* macrophages were found to be linked with cell-adhesion-related pathways, phagocytic signaling routes, and the GTPase cycle, processes previously linked to this state. This highlighted the capability of the MONET decomposition tool in extracting subnetworks associated with the most important signaling routes in different macrophage phenotypes.

#### 3. NOODAI workflow

The webtool, hosted at omics-oracle.com, features a comprehensive user manual that explains input parameters and provides guidance on general usage. The platform relies on DigitalOcean cloud service infrastructure and incorporates essential security configurations to ensure the protection of the uploaded data. A cloud solution was chosen as it is easily scalable. The interface was designed in R Shiny (Chang, et al., 2021) and it makes use of *uuid* (Urbanek and Ts'o, 2022), *shinyjs* (Attali, 2021), *bslib* (Sievert, et al., 2023) and *shinyWidgets* packages (Perrier, et al., 2023) as well as *parallel* (R Core Team, 2022) and *callr* (Csárdi and Chang, 2022) that make possible the dynamic use of the web service. For each client submission, a background process is started which allows users to close their browsers while they wait for the results. The omics-oracle interfaces and user manuals were designed in HTML and CSS with bootstrap v5. The source code of the NOODAI pipeline is available on GitHub (Totu, 2024). NOODAI relies on *ggplot2* (Wickham, 2020) for graphical design and *biomaRt* (Durinck, et al., 2009) for mapping UniProt IDs to NCBI IDs when species other than those for which the pre-loaded datasets are available are analyzed. Additional packages that are used during the analysis are *devtools* (Wickham, et al., 2022), *reshape* (Wickham, 2007), *moments* (Komsta and Novomestky, 2022), *EnvStats* (Millard, 2013), *readr* (Wickham, et al., 2023), *readxl* (Wickham, et al., 2023), *openxlsx* (Schauberger and Walker, 2023), *tidygraph* (Pedersen, 2023), *ggpubr* (Kassambara, 2023), *gtools* (Bolker and Warnes, 2022), *stringr* (Wickham and Wickham, 2019), *ggcorrplot* (Kassambara, 2019), *centiserve* (Jalili, et al., 2015), *RColorBrewer* (Neuwirth and Brewer, 2014), *gridBase* (Murrell, 2014), *circlize* (Gu, et al., 2014), *clusterProfiler* (Wu, et al., 2021), *igraph* (Csardi and Nepusz, 2006)*, dplyr* (Wickham, et al., 2023) and *CINNA (Ashtiani, et al., 2019)*. The webtool is released under GPL license v3.0.

The NOODAI webtool workflow consists of two main parts: data upload together with the selection of input parameters and the subsequent results download. To access the results after the analyses are finished, the user navigates to the "Results download" tab. Here, the files can be downloaded as a compressed archive using the assigned token ID generated upon submission. The results of an analysis are stored on the server for a maximum of 5 days, and the associated token IDs are randomly generated to ensure data security.

Each of the 4 sections of the analysis pipeline can be considered as an independent Segment of the workflow that can be individually accessed through the platform (Network construction and node centrality – Network decomposition (MONET) – Module signaling pathways analysis – Summary plots and report generation). The main interface (Figure S2A) allows users to efficiently run the entire analysis pipeline (all 4 Segments) requiring only a few inputs. Each field has an associated description. Most fields have default values, while some require mandatory input from the user. The fields marked in Figure S2A are described as follows:

1. The left Demo button allows automatic upload of the demo dataset available on the platform with pre-configured input parameters. Pressing the Submit button starts the analysis of the demo dataset.
2. In the filed "Conditions Names", users must upload the names of each analyzed phenotype or condition separated by commas. Quotes and underscores should not be used. For instance, for the demo dataset this field should be "M1,M2a,M2c".
3. Under "Conditions Contrasts", users are required to upload the comparison groups. In the case of the demo dataset, a comparison group is represented by the elements upregulated in one phenotype/condition compared to another, and the input for this field has the following structure:

   "M1vsM2a,M2avsM1,M1vsM2c,M2cvsM1,M2avsM2c,M2cvsM2a". Each contrast is composed of Condition Names joined by "vs" and must be identical to the Excel sheets names of the uploaded omics files. Different contrasts must be separated by commas.
4. In the "Omics Files Archive" field users are required to upload a single archive which contains one Excel file for each omics layer. Each Excel file must include at least one sheet. As mentioned above, the names of each sheet should match precisely the comparison groups provided in the "Sample Contrasts" field. In the case of the demo dataset, an Excel file for each omics layer contains 6 sheets with the following names "M1vsM2a", "M2avsM1", "M1vsM2c", "M2cvsM1", "M2avsM2c" and "M2cvsM2a". The Excel sheets must include a column named "UniProt_ChEBI". This column should have only UniProt or ChEBI IDs.
5. The default "MONET method" string parameters are for the MONET M1 method. Users have the option to change the method to R1 or K1. The avgk parameter that influences the average module size can also be adjusted as needed. The format must follow the one in the default setting.
6. Pathway over-representation analysis computed in the 4$^{th}$ Segment can be performed with multiple signaling pathway databases, including Reactome, Wikipathways, BioCarta, PID, NetPath, HumanCyc, INOH, and SMPDB. Users are responsible for complying with the licensing terms of these databases.
7. The default "BioMart dataset" field is pre-set for mapping human protein IDs, but it can be adapted to other species. The IDs are mapped with the biomaRt R package.
8. A user can choose to use pre-formatted interaction files available on the server or add the new PPI dataset by clicking on the blue plus sign. To use the webtool for organisms other than the 13 for which the interaction datasets are pre-loaded or

to use other types of interactions, it is mandatory to upload a custom pre-formatted interaction file.

9. The optional “Email address” field allows users to receive an email notification when the analysis is completed.

When the “Submit” button is pressed, the analysis starts and a unique token is generated. This token should be used in the “Results download” tab for retrieving the final results when the analysis is finished.

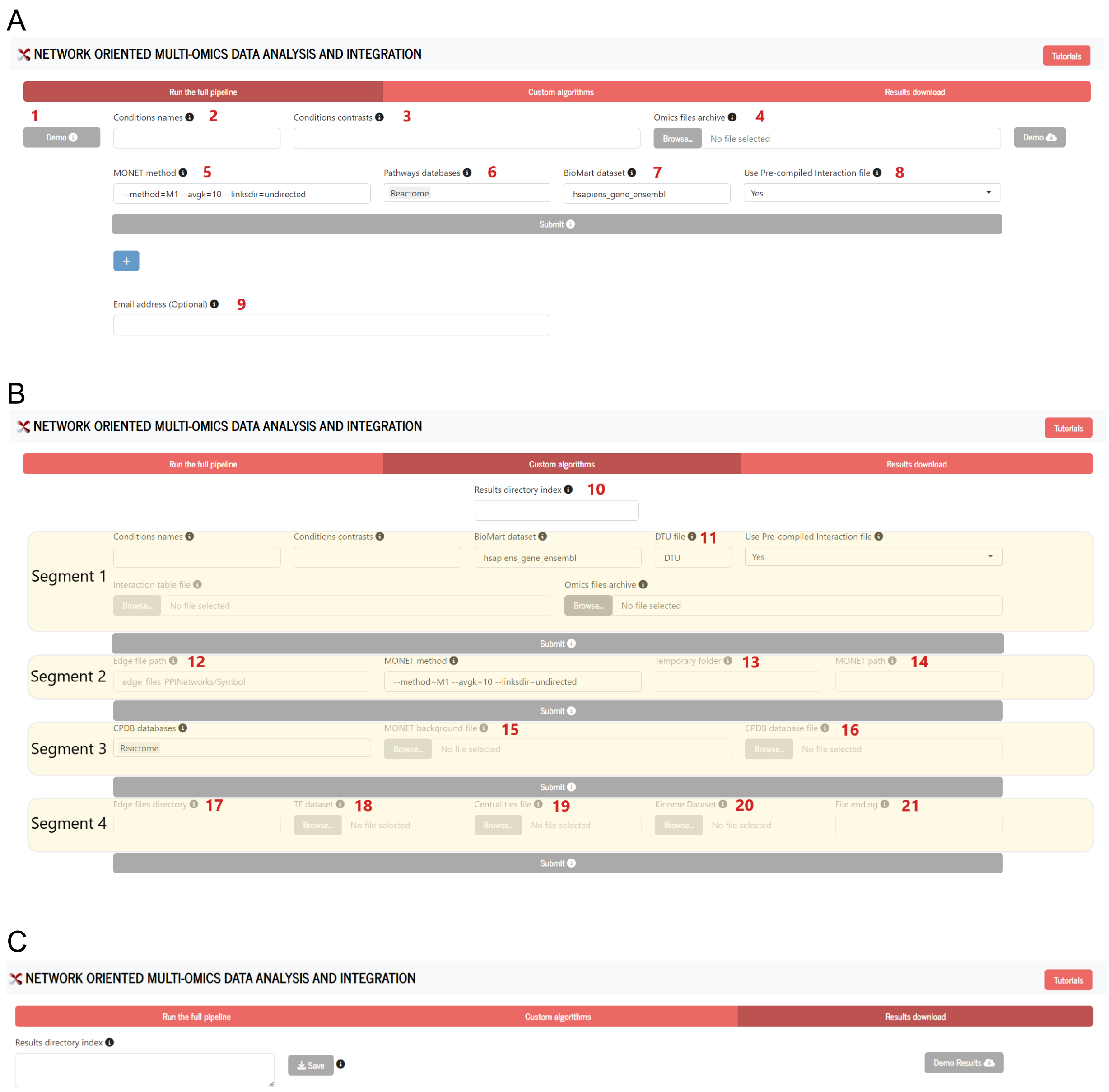

**Figure S2.** NOODAI interface with the numbering of the fields based on the order of appearance: A) Complete pipeline run B) Individual algorithms tab and C) Results download panel.

In addition to running the entire analysis pipeline in a single run, the NOODAI interface allows users to customize the parameters of each individual analysis segment (Figure S2B) and run the segment separately. However, this can be done after the complete analysis pipeline is finalized from the “Run the full pipeline” tab and a token is assigned to it. The following description applies to the fields within the “Custom Algorithms” tab that support optimization of parameters for individual segments:

10. An analysis token generated for the execution of the full analysis pipeline can be used in the "Results directory index" for running an analysis segment independently.
11. If the input omics datasets include splicing data please enter the exact Excel file name (without extension) into the "DTU file" field.
12. By default, the names reported for the nodes in the MONET modules and in the results of the pathway analysis are official gene names. They are stored as Gene Names in the "Symbol" folder on the server. To use UniProt IDs instead of Gene names, users can use the "Edge file path" field and change there the folder path from “Symbol” to “Uniprot”.
13. - 14. These two fields are relevant when the pipeline is downloaded from GitHub and used locally. The webtool was designed to be run locally as well. For this, it is necessary to create a temporary folder in which intermediate files and results from the MONET analysis are stored. The name of the folder should be defined in the “Temporary folder” field. The temporary files created in this folder are automatically deleted at the end of the data processing. In addition, when NOODAI is run locally, the MONET executable path must be provided in the “MONET path” field. There is also a possibility for the users to use a different background list for the overrepresentation analysis (default are all entities from the joint PPI network). An alternative background file can be uploaded using the "MONET background file" field.
16. The pathways knowledge-based datasets used for the over-representation analysis are extracted from the CPDB database. The user can change and upload their own pathway annotations by using the “CPDB database file” field and following the same formatting scheme.
17. By default, circular plots and the final summary report use official gene names. Users preferring to use UniProt IDs can switch to the “Uniprot” folder as mentioned in point 12 above. The "Edge files directory" field must match the "Edge file path".

Therefore, both fields should be formatted identically when changing the edge folder.

18. A different dataset with known transcription factors can be uploaded under the “TF database” field. The pre-loaded TFs are from AnimalTFDB (Shen, et al., 2023). If users would like to highlight other elements and not transcription factors in the circular diagrams, they can use this field to provide their own entries which will be visualized with their interaction partners when they are ranked as highly central.
19. NOODAI relies on a PPI network constructed by concatenating the networks from individual omics profiles. Results output files also include centrality metrics for individual omics layers. If the users wish to generate the summary report and plots for a specific omics input dataset, they can upload the respective centrality file under the "Centralities file" field. If one would like to have the summary report and plots for a different metric than the current-flow betweenness centrality, the file can be modified offline and uploaded again on the platform.
20. The user can choose to use a different list of kinases instead of the default one available on the server. This can be uploaded under the "Kinome Database" field. This may be also used if the user wants to replace kinases with phosphatases, epigenetic regulators or other class of proteins.
21. By default, the "File ending" field is set to "Total", indicating that the network built from all omics layers is analyzed for the final plots and report. In order to generate plots only for a certain omics dataset, another centrality file should be used, as described in point 19 above, and the “File ending” field should be updated with the name of the input omics file name.

The "Results download" tab (Figure S2C) allows users to download the results generated from the analysis of the demo dataset or to retrieve the results from their own analysis in the form of an archive file.

The platform offers users flexibility with regard to the type of data that can be analyzed and the parameters used for the analysis. Its default inputs are protein and small molecule IDs and it has pre-loaded datasets for thirteen species, but the pipeline can be used for any other species and omics features as well. For this, users must provide the correct BioMart dataset name, a pre-formatted feature-feature interaction file, a reference signaling pathway database, and a custom TF dataset. For the pre-loaded species, only the BioMart dataset name must be properly set up. For more details, the users are encouraged to consult the online documentation available on the platform.
